# Cyanolichen microbiome contains novel viruses that encode genes to promote microbial metabolism

**DOI:** 10.1101/2021.06.15.448434

**Authors:** Alise J. Ponsero, Bonnie L. Hurwitz, Nicolas Magain, Jolanta Miadlikowska, François Lutzoni, Jana M. U’Ren

## Abstract

Lichen thalli are formed through the symbiotic association of a filamentous fungus and photosynthetic green alga and/or cyanobacterium. Recent studies have revealed lichens also host highly diverse communities of secondary fungal and bacterial symbionts, yet few studies have examined the viral component within these complex symbioses. Here, we describe viral biodiversity and functions in cyanolichens collected from across North America and Europe. As current machine learning viral-detection tools are not trained on complex eukaryotic metagenomes, we first developed efficient methods to remove eukaryotic reads prior to viral detection and a custom pipeline to validate viral contigs predicted with three machine-learning methods. Our resulting high-quality viral data illustrate that every cyanolichen thallus contains diverse viruses that are distinct from viruses in other terrestrial ecosystems. In addition to cyanobacteria, predicted viral hosts include other lichen-associated bacterial lineages and algae, although a large fraction of viral contigs had no host prediction. Functional annotation of cyanolichen viral sequences reveals numerous viral-encoded auxiliary metabolic genes (AMGs) involved in amino acid, nucleotide, and carbohydrate metabolism, including AMGs for secondary metabolism (antibiotics and antimicrobials) and fatty acid biosynthesis. Overall, the diversity of cyanolichen AMGs suggests that viruses may alter microbial interactions within these complex symbiotic assemblages.

## Main Text

Lichens—defined as the symbiotic association between a filamentous fungus (mycobiont) and at least one photosynthetic organism (photobiont)—grow on a broad array of substrates in terrestrial, freshwater, and marine intertidal ecosystems from the poles to the tropics [1]. The vast majority of lichens contain green alga (Chlorophyta) as their main photobiont, but >1,500 species of lichens have *Nostoc* cyanobacteria either as primary photobionts (forming bi-membered lichens) or secondary photobionts (forming tri-membered lichens with a green alga as the primary photobiont) [2]. The cyanobiont provides photosynthate and fixed nitrogen, while the fungal partner provides the photobiont with protection from light, carbon dioxide, and inorganic ions [2]. Recent molecular studies illustrate that lichens also contain cryptic secondary bacterial and fungal symbionts [3–6], yet few studies to date have examined viruses that associate with these complex microbial communities (but see [7]).

Viruses infect all domains of life and are the most abundant biological entity on Earth [8]. Previous studies have shown that viruses that infect bacteria are evolutionarily tuned to their hosts in a given environment toward two main goals: promote viral replication for lysis or co-exist within the host genome as a prophage [9]. To do so, viruses encode host genes (i.e., auxiliary metabolic genes (AMGs; [10]) that promote viral replication through manipulation of the host’s metabolism. For example, marine cyanophages encode *psbA* to increase host photosynthesis and drive replication when host *psbA* is inhibited by high light [11]. Viral AMGs can provide important clues into host adaptation, metabolic bottlenecks, key ecosystem functions, and interactions among members of a microbial community [9, 12].

Viral sequences in metagenomes can be detected using reference-based methods, but these methods often are hampered by the limited diversity of viral genomes in reference databases (see [13]). Thus, an emerging approach is to apply machine learning (ML) algorithms that use composition-based pattern detection. ML models identify a set of features that signal a viral origin (e.g., relative synonymous codon usage, gene density, strand shifts, and the number of significant hits against the pVOGs database), thus generalizing the identification of all viral sequences and enabling better detection of novel viruses [13–15]. These new approaches provide exciting avenues for detecting novel viral sequences in metagenomes and are key to investigating complex microbial symbioses such as lichens.

Here, we explored viral biodiversity in 11 cyanolichen metagenomes (Table S1) representing nine species of the genus *Peltigera* sampled from North America, Finland, Iceland, and Panama, and one species from the sister genus *Solorina* (Fig. 1a) [16]. We predicted viruses with three ML tools (MARVEL, Vibrant, VirSorter) [13–15]. However, as repetitive regions of eukaryotic genomes can be falsely identified as viral [17], we first developed a pipeline to remove eukaryotic sequences prior to viral detection and applied a stringent cutoff to limit false positives (i.e., minimum 10% of ORFs/contig hit a previously identified viral protein) (Fig. S1).

**Figure 1.**
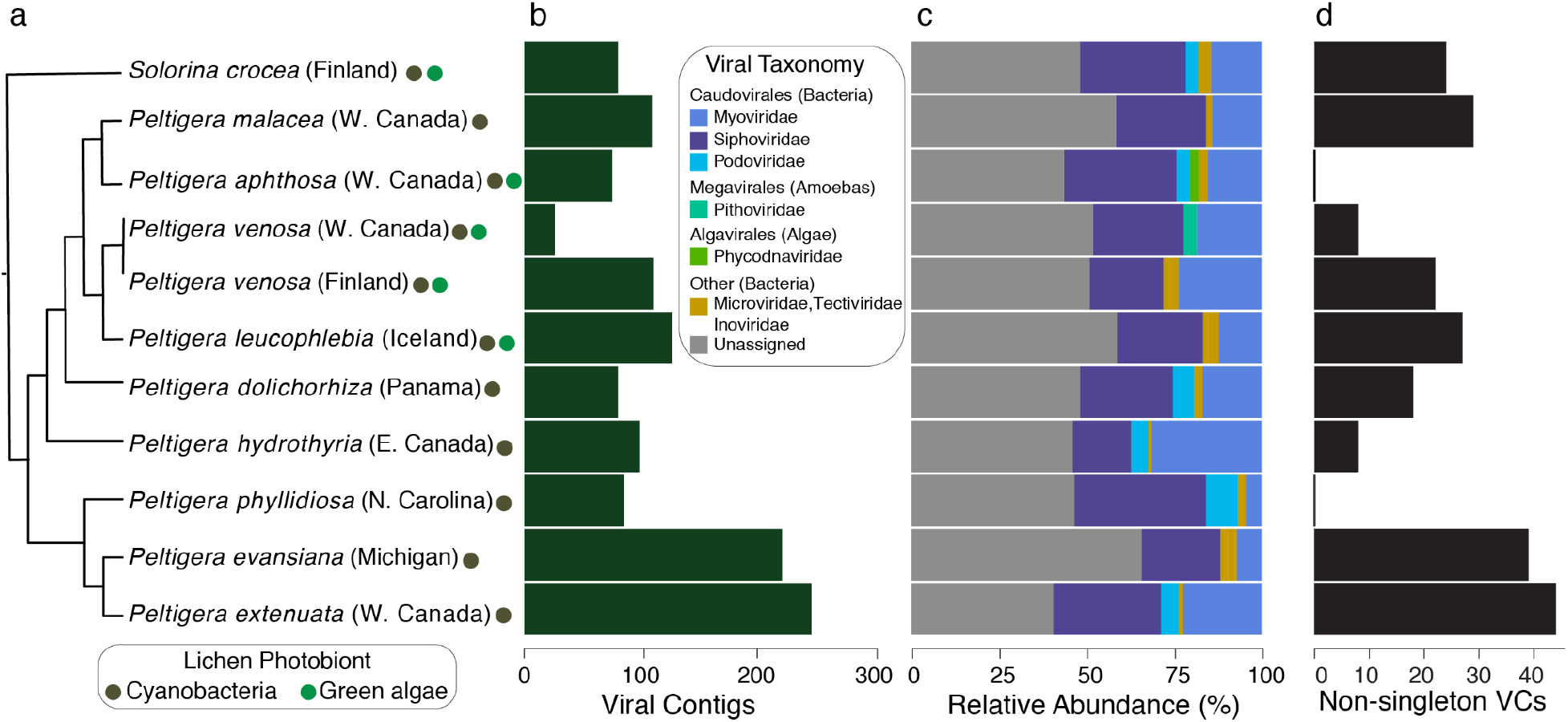
Cyanolichens harbor diverse viral communities that infect prokaryotic and eukaryotic hosts in lichen thalli. (**a**) Schematic tree illustrating the phylogenetic relationships among lichen mycobionts (Peltigerales) (see Supplementary Materials). Colored circles indicate the photobiont composition of each lichen (see legend). Names associated with two colored circles are tri-membered lichens, whereas names associated with one colored circle are bi-membered cyanolichens. (**b**) Bar graph of the number of cyanolichen contigs predicted per sample, which accounted for 0.6 to 3.4% of the total reads per sample. (**c**) Taxonomic classification of viral contigs identified 15 viral families that infect prokaryotic (n = 802 contigs) and eukaryotic hosts (n = 28 contigs), although 471 (36.2%) contigs could not be classified. (**d**) Distribution of non-singleton viral clusters (VCs) among samples. Among the 133 VCs, 50 occurred only in a single cyanolichen sample.

Every cyanolichen thallus contained viral sequences (range 27 to 254; Fig. 1b), although we observed no pattern between viral abundance and thallus type (bi- or tri-membered) or geographic location. In total, we predicted 1,301 non-redundant viral contigs (including 116 predicted prophage sequences and 27 complete viral genomes) with high confidence across all metagenomes. The majority of viral contigs were classified as bacteriophages from the *Caudovirales* (61.2%) (Fig. 1c). We detected few prophages, suggesting that bacteriophages may be predominantly lytic. Twenty-eight contigs also matched nine viral families that infect eukaryotes, including viruses of *Phycodnaviridae* that infect green algae [18] (Fig. 1c). Our taxonomic results are consistent with the observed bacterial communities in these samples (dominated by Proteobacteria and Cyanobacteria; Fig. S2a), as well as our *in silico* predictions of viral hosts (Fig. S2b). Additional eukaryotic viruses likely associate with algal photobionts of tri-membered lichens (see Fig. 1a) and fungal mycobionts in lichen thalli [7], but biases in both computational methods and databases towards phages may have limited their detection and classification (e.g., 36.2% of viruses were unclassified; Fig. 1c). Additionally, many algal and fungal viruses are dsRNA viruses that cannot be detected with metagenomic data [7, 19].

To assess the novelty of cyanolichen viruses, we compared our contigs to >400,000 previously published terrestrial viral contigs and genomes. Few of our cyanolichen contigs form viral clusters (VCs) with non-lichen sequences (Fig. S3) and the majority (n = 966 contigs) were classified as singletons or outlier VCs, highlighting the novelty of cyanolichen viral sequences. The remaining 335 contigs represent 133 VCs across the 11 metagenomes (Fig. 1d). These non-singleton VCs had limited overlap to previously published sequences (105 VCs contained sequences only from lichens) or among different cyanolichen samples (Fig. S4). Increasing sequencing depth would likely recover additional viruses, but the lack of VC overlap among cyanolichen samples also may reflect high geographic turnover of non-cyanobacterial lichen-associated bacterial communities [3] similar to turnover of marine viral communities according to host identity and physical or chemical properties of the environment [12, 20].

To assess the functional diversity of cyanolichen viruses we predicted open reading frames (ORFs) (n = 21,855), which we then clustered into 20,776 protein clusters (PCs). Overall, >19,000 PCs were singletons (96%), illustrating little functional overlap among viruses and samples, consistent with highly novel phage reported in other terrestrial ecosystems [8].

Functional annotation of cyanolichen viral genes identified 550 ORFs (from 249 contigs, including circular genomes) with significant matches to metabolic KEGG HMM profiles (Fig. 2). AMGs in cyanolichens include KEGG pathways for amino acid, nucleotide, secondary metabolite (e.g., antibiotics), and carbohydrate metabolism (e.g., *psbA* [12]) (Fig. 2; Tables S2-S3), consistent with the ability of phages in other ecosystems to encode host genes to drive host metabolism [10–12].

**Figure 2.**
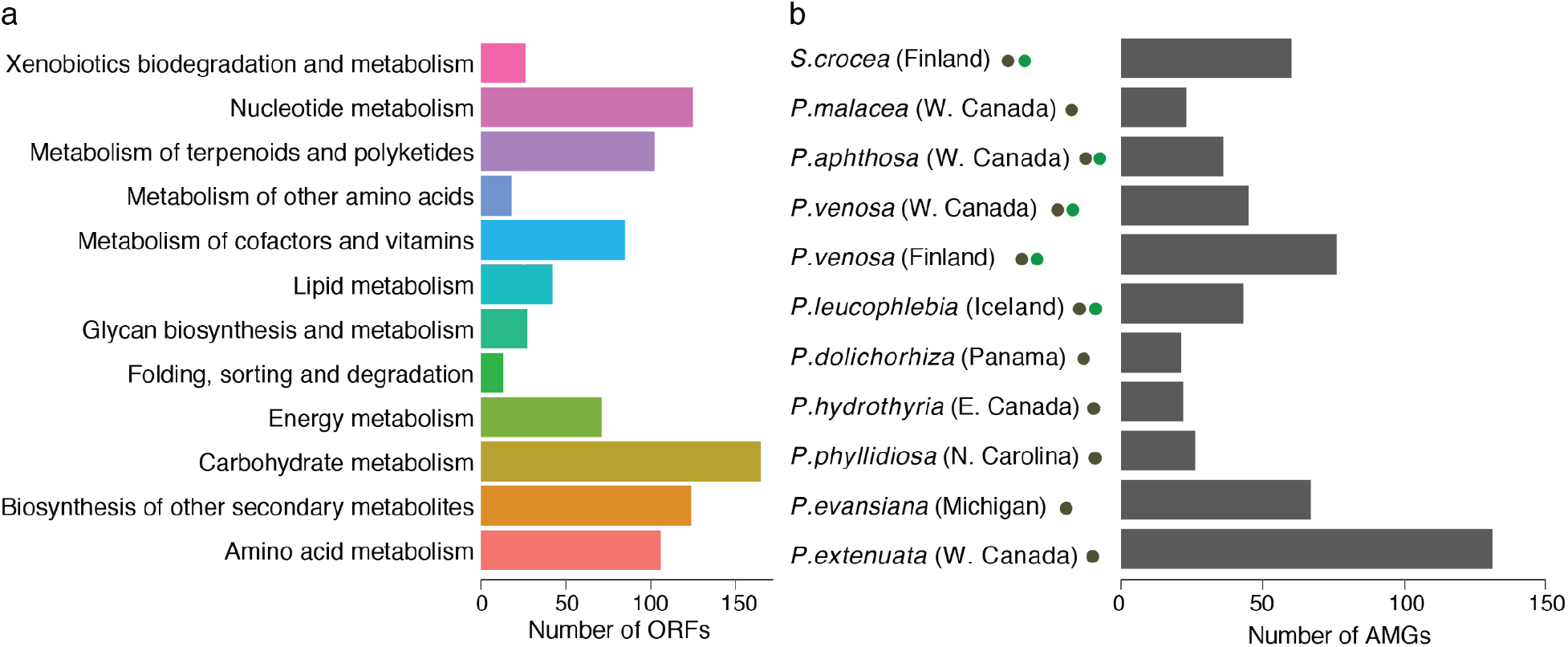
Cyanolichen viruses encode diverse auxiliary metabolic genes (AMGs) for amino acids, nucleotides, carbohydrates, and secondary metabolite metabolism. (**a**) Number of AMGs per KEGG pathway. AMGs were found on 250 contigs, including 9 of the 27 circular viral genomes; (**b**) Number of AMGs per cyanolichen sample.

In conclusion, a large variety of tools and approaches have been used to identify viruses in marine, human-associated, and soil metagenomes, yet viral detection remains challenging in complex and under-explored eukaryotic host-associated metagenomes. Here, we illustrate the diversity, novelty, and functional potential of viruses in cyanolichens and identify AMGs for metabolic pathways not previously described in viruses in other ecosystems (e.g., dTDP-L-rhamnose biosynthesis, fatty acid biosynthesis). The abundance and diversity of viral AMGs in cyanolichens, including numerous AMGs for secondary metabolism, suggests viruses may modify interactions among complex microbial partners within lichens. Although viral novelty and microbial complexity in cyanolichen metagenomes limited our ability to link AMGs to specific hosts, future work will seek to decipher these phage-host associations and the functional roles of AMGs in the lichen symbiosis.

## Acknowledgments

Funding for this work was provided by NSF [OCE-1639614 Planet Microbe to BLH; DEB‐ 1556995 and DEB-1541548 to FL and JM] and Gordon and Betty Moore Foundation [GBMF 8751 to BLH]. Lichen sampling was funded by NSF grants DEB-1046065 and DEB-1541548 to FL and JM.

## Author Contributions

Designed research: JMU, BLH; Performed research: AJP, JMU; Contributed data or analytic tools: NM, FL, JM; Analyzed data: AJP, JMU, BLH; Wrote the paper: AJP, JMU, BLH, with contributions from all authors.

## Availability of Data and Code

Data used in this study are available at the National Center for Biotechnology Information (NCBI) Sequence Read Archive (SRA) (see Table S1 for accession numbers). All code is available on GitHub (https://github.com/aponsero/) including the complete pipeline for (i) identifying viral contigs using VirSorter, Vibrant, and MARVEL (Viral_hunt_snakemake); (ii) validating that contigs are of viral origin (Viral_confirmation_snakemake); and (iii) code to run VirFinder in parallel on a High-Performance Computer (HPC). Viral reads are available on figshare (10.6084/m9.figshare.c.5444376).

