## Supplementary Materials for "Cyanolichen microbiome contains novel viruses that encode genes to promote microbial metabolism"

**This file includes:**

Materials and Methods

Supplementary Tables S1-S3

Supplementary Figures S1-S6

### Materials and Methods

***Sampling, Metagenomic Sequencing, Quality Control and Assembly.*** To investigate the diversity, composition, and function of viral communities associated with cyanolichen thalli, we analyzed metagenomic data from eleven cyanolichens previously collected from montane and boreal forests in North America and Europe (Table S1) [16]. Lichen mycobionts span the phylogenetic diversity of Peltigerales, including nine species of *Peltigera* and one species of the sister genus *Solorina* (Peltigeraceae, Peltigerales) [16]. Phylogenetic relationships among lichen mycobionts (e.g., Fig. 1 and elsewhere) are based on a phylogeny of the genus *Peltigera* based on ref [21] that is available in T-BAS (https://tbas.hpc.ncsu.edu/tbas) [22, 23].

DNA library preparation (DNA-Seq, 300 bp insert) and metagenomic sequencing (Illumina HiSeq 4000 150-bp PE) was performed by the Sequencing and Genomic Technologies Shared Resource, Duke Center for Genomic and Computational Biology, Duke University (Durham, North Carolina, USA). We trimmed and filtered raw reads from each cyanolichen metagenome using TrimGalore! V0.4.5 [24]. Low-quality base calls (quality score below 20 Phred) were removed from the 3' end of the reads, as well as adapters and poly-G sequences. After trimming, we removed reads <20 bp in length. Quality-filtered reads were assembled using Megahit v1.2.9 using the default parameters [25, 26]. As lichens are complex microbial communities, the resulting reads for each metagenome represent eukaryotic and prokaryotic microbes, including fungi (mycobiont [1] and endolichenic fungi: [4]), bacteria (cyanobacterial photobiont [27] and secondary epiphytic and endophytic bacteria [3]), as well as green algal photobionts (tri-membered only [1], i.e., one mycobiont species and two photobiont species; see Table S1).

***Evaluation of ML tools for detecting viruses in eukaryotic dominated metagenomes.*** To date, machine learning (ML) tools for detecting viruses have been primarily evaluated on metagenomes dominated by prokaryotic microbes without eukaryotic sequences. However, previous analyses have demonstrated the high rate of viral false positives that can result from eukaryotic sequences for some computational approaches [17], thus we first evaluated the robustness of computational tools to detect viruses using a simulated mock community representing lichen metagenomes that contains fungal, algal, cyanobacterial (*Nostoc* sp.), and cyanophage genomes (see Table S4 for list of genomes).

Briefly, genome sequences were trimmed to 500, 1000, 3000 and 5000 bp fragments. For each fragment length, the sequences were then randomly selected to create 10 replicate evaluation sets, each composed of 1,000 sequences of fungal, algal, bacterial and viral sequences (4,000 total sequences per set). Each replicate mock community was analyzed with VirSorter, VirFinder, and Vibrant [13, 15, 28]. VirFinder v1.1 was run on the evaluation sets using the default model. A hit was considered as viral for a score equal or above 0.96. VirSorter v1.0.5 was run on the evaluation sets using the default database. All hit categories were taken into account. Vibrant v1.0.1 was run on the evaluation sets using the default database. All hit categories were taken into account.

Unlike VirSorter, VirFinder, and Vibrant that directly analyze contigs, MARVEL [14] requires a cluster step prior to analysis to bin contigs that are likely from the same organisms. Thus, to evaluate MARVEL, we generated artificial bins containing 10 sequences from the same genome. Each test, corresponding to a specific fragment length, was performed in ten randomly sampled replicates of 100 bins. Bins were submitted to MARVEL and predictions were evaluated for true positive rates, specificity, accuracy, and F1 score according to standard formulae. MARVEL v0.2 was run on the evaluation bins using the default settings. All bins retrieved by MARVEL were considered in the final comparison of tools. Overall, we found that VirFinder had the highest viral false positive rate when tested with eukaryotic sequences (Fig. S5), which we attribute to the absence of eukaryotic sequences in the training set for the VirFinder model (also see [17]). As such, we assessed whether including fungi and micro-eukaryotic sequences in the VirFinder training set would improve the tool's performance by developing three new models trained using complete fungi and algal genomes (i.e., VirFinder A,B,C; see Data Availability and Code). The revised VirFinder models were trained using 15,000 sequences from RefSeq Viral genomes, 5,000 sequences from Ascomycota genomes, 5,000 sequences from *Nostoc* genomes, and 5,000 sequences from green algal genomes. The genomes were retrieved from the RefSeq [29] and PATRIC [30] databases and sequences were randomly fragmented into lengths of 5,000 bp. We created three random subsets of these sequences to train three independent models (VirFinder A,B,C) and evaluated their performance using the same mock community and methods as above. Although the new models showed a lower false positive rate for algal and fungal sequences (Fig. S5), the number of false positives was still higher than observed for VirSorter, MARVEL and Vibrant. As a result, we excluded VirFinder from the set of tools we used to identify viruses in cyanolichen metagenomes**.**

***Identification of viral sequences in cyanolichen metagenomes.*** After evaluating computational tools for detecting viruses using mock community datasets and confirming low rates of viral false positives, we used VirSorter, Vibrant, and MARVEL [13–15] to identify viral sequences in cyanolichen metagenomes (Fig. S1). All programs were run using the default settings. MARVEL v0.2 was run on the assembled metagenomes after first binning using Metabat2 v2.12.1 [31]. For each putative viral contig identified, we implemented a validation pipeline to further confirm viral origins of putative contigs (Fig. S1, See Data Availability and Code).

Briefly, the validation pipeline confirmed putative viral contigs using the following methods. First, low coverage contigs were removed by mapping the raw reads (after QC) against the assembled contigs using Bowtie2 [32] and BBMap/samtools [33, 34]. Contigs with an average coverage depth less than 5x were removed. Second, we used Prodigal v2.6.3 [35] using default settings for gene calling on each putative viral contig. The predicted proteins were compared to pVOGs (downloaded on Jan. 30, 2020) [36] and PVF from the IMG/VR v2.0 database [37] (downloaded on Jan. 30, 2020) profiles using HMMer v3.3 [23]. Sequence alignments were considered significant if the e-value was ≤0.01. We considered the viral contig to be validated if at least three ORFs and a minimum of 10% of all ORFs were annotated as being viral. Putative prophage sequences were considered viral if the contigs contained at least three ORFs with a viral annotation. Finally, potential host and sequencing contamination was removed by comparing contigs to fungal genomes from the RefSeq database (downloaded on June 2019) and the PhiX genome using BLASTp [38]. We discarded viral contigs with a BLASTp e-value <0.01 and a hit length >500bp. The complete pipeline for validating that contigs are of viral origin is available as a snakemake workflow (see Data Availability and Code).

After the validation pipeline, several viral contigs appeared to represent circular viral genomes (Table S5a). We confirmed the circularity of non-prophage contigs by identifying matches between the start and end of the contig. Briefly, 100 bp at the start and end of each contig was aligned using BLASTn [38]. Potential circular contigs were defined by a hit longer than 45 bp between the 100 bp fragment at the beginning and end with a percent identity >85%. Overall, we predicted 1,301 high confidence viral contigs, with 27 contigs showing a circularity that is consistent with complete viral genomes. Vibrant retrieved the largest number of putative viral contigs (n = 962), followed by VirSorter (n = 598), and then MARVEL (n = 136) (Fig. S6a). Comparison of the viral contigs predicted by each program indicated that each method predicted distinct viral sequences, illustrating that use of multiple ML approaches allows for broader retrieval of putative viral sequences. We attributed differences among ML methods to the different parameters used by each method. For example, Vibrant viral detection permits the inclusion of shorter viral sequences, whereas Virsorter tends to retrieve longer sequences (Fig. S6b). In total, only 59 viral contigs were identified by all three tools (Fig. S6a).

Overall, these viral detection methods retrieved very few contigs with prophage (9% of viral contigs). To assess whether this result was a biological phenomenon or methodological artifact (i.e., difficulty in retrieving prophages from assembled contigs or limitations in the length and quality of the assembly), we searched for prophage in publicly available lichen-associated *Nostoc* genomes retrieved from NCBI (genome ID: SAMN03354403, SAMN07192327, SAMN03445855, SAMN07191894, SAMN07191895, SAMN07191897). We used VirSorter v1.0.5 (default database [13]) to predict prophages on scaffolds of six *Nostoc* genomes. We found 54 partial prophage sequences (VirSorter categories 2 and 3); however, no intact prophage sequence could be detected in these genomes (Table S8). Previous studies show a reduction in genome size and proliferation of pseudogenes in lichen-associated *Nostoc* compared to free-living species [39], which could also lead to a degradation of prophages integrated into *Nostoc* genomes and explain the absence of complete integrated prophages here.

***Identification of cyanolichen viral taxonomy and putative bacterial hosts***. As lichens represent complex communities with both primary cyanobacterial symbionts in the genus *Nostoc*, as well as secondary bacterial symbionts that occur on the surface and interiors of thalli, we classified the taxonomy of viral contigs with DemoVir [40] using the Trembl database with non-redundant sequences at 95% (https://doi.org/10.6084/m9.figshare.5822166.v1). Overall, 802 (61.6%) of the retrieved viral contigs were classified as bacteriophages (Table S6). The majority of phages were classified in the order *Caudovirales* (n = 796 contigs of *Siphoviridae*, *Myoviridae*, *Podoviridae*, and unclassified *Caudovirales*), although six contigs were classified as phages in the families *Microviridae*, *Tectiviridae,* and *Inoviridae* (Table S6). Only 2.2% of the retrieved contigs were classified as eukaryotic viruses (n = 28). Putative eukaryotic viruses had matches to nine viral families, including the viral families *Caulimoviridae* and *Phycodnaviridae* whose hosts include plants and green algae, respectively (Table S6). The remaining 36.2% (n = 471) of contigs could not be classified (Table S6).

Next, we used the PHIS detector to predict bacteriophage hosts *in silico [41]*. The resulting GenBank accession numbers for each putative host were compared to the NCBI taxonomy database to obtain consensus taxonomic classification (Fig. S2b) with a Least Common Ancestor (LCA) approach (sensu MEGAN; [42]). In cases where predicted taxonomic assignments were incongruent (e.g., predicted hosts represent two different bacterial phyla), hosts were categorized as ‘undecided’. In total, hosts were predicted for ~25% (n = 324) of the bacteriophage contigs from cyanolichens (Table S6). The most common identifiable hosts across the cyanolichen viral contigs belonged to the bacterial classes Alphaproteobacteria (140 contigs), Nostocales (36 contigs), and Betaproteobacteria (13 contigs) (Fig. S2b). Among the 27 circular genomes retrieved, only five could be associated with a potential host (one host predicted as Gemmatimonadetes, one betaproteobacteria, and three phages of *Nostoc*).

We then compared the predicted bacterial host population to the observed bacterial communities in cyanolichen thalli (Fig. S2a). We classified all unassembled metagenomic reads using Centrifuge version 1.0.3 [43] (default settings) after QC filtering against the “p_compressed+h+v” compressed database (released April 15, 2018). Overall, predicted viral hosts are consistent with the observed taxonomy of the cyanolichen bacterial communities. For example, Proteobacteria and Cyanobacteria are the most abundant bacterial phyla present in the total metagenomes of bi-membered cyanolichen samples (Fig. S2a). Discrepancies between observed bacteria taxonomy and predicted bacteriophage hosts are likely due to the large fraction of bacteriophage with unknown host predictions (Fig. S2b).

Differences in the relative abundance of cyanobacteria among metagenomes (Fig. S2a) are likely the result of structural differences between bi- and tri-membered cyanolichens. For example, we observed fewer cyanobacterial reads in the tri-membered lichens *P. venosa* and *P. leucophlebia*, where the primary photobionts are green algae and cyanobacterial symbionts are restricted to cephalodia. In contrast, bi-membered cyanolichens that do not contain green algal photobionts generally contained a higher proportion of cyanobacterial reads (Fig. S2a). Differences in the relative abundance of *Nostoc* among samples is likely to impact the composition of viral sequences identified in each metagenome.

***Comparisons of cyanolichen viral contigs to other terrestrial ecosystems.*** To assess the novelty of viruses found within cyanolichens, we compared predicted cyanolichen viral contigs to viral contigs from other terrestrial ecosystems. Briefly, we downloaded 466,091 non-singleton vOTU from "Soil", "Rhizoplane","Rhizosphere", "Phyllosphere" and "Plant litter" ecosystem type from the the IMG/VR database (version 4 - 2018-07-01_4) [37], as well as 2,191 reference phage genomes from NCBI Viral RefSeq version 88 [44]. All contigs were then annotated using Prodigal v2.6.3 and genus-level “viral clusters” (VCs) were computed with vConTACT 2.0 [45].

We observed little overlap of VCs among cyanolichens (Fig. S4). A network diagram of clusters illustrates that very few VCs shared between cyanolichens and other terrestrial ecosystems (Fig. S3). The majority of cyanolichen viral contigs (n = 966) were classified as singletons or outlier VCs and the remaining 335 contigs represent 133 VCs across the 11 metagenomes (Fig. 1d). These non-singleton VCs had limited overlap to reference sequences (105 VCs only from lichens). Only 230 contigs clustered with previously published viral sequences from NCBI RefSeq or terrestrial viral contigs published in IMG/VR.

***Clustering of cyanolichen viral proteins***. Putative proteins obtained by Prodigal were clustered using CD-HIT version 4.8.1 (built on Oct 26, 2019) [46], using a sequence identity threshold of 60% identity and 80% coverage [47]. Pfam annotations for the protein clusters were obtained using the HMMERv3.3 and the Pfam HMM profiles v33.1 (May 2020) [48, 49]. The cutoff e-value was set to 0.01. We further confirmed the novelty of viruses among cyanolichen samples by investigating the shared protein clusters among samples. Prodigal identified 21,855 open reading frames (ORFs) across all cyanolichen viral contigs. We then clustered putative proteins into protein clusters (PCs) with a sequence identity threshold of 60% identity and 80% coverage, which yielded a total of 20,776 PCs, 33% of which were annotated with Pfam, which is higher than what was reported for soil viromes studies (10%) [50]. However, only 4% of PCs (n = 800) contained more than one sequence (i.e., non-singleton).

***Detection of AMGs in cyanolichen viruses.*** Many phages have been shown to encode metabolic genes of host origin which may be retained in the viral genome, called Auxiliary Metabolic Genes (AMG). AMGs are thought to enhance viral replication by bolstering the metabolism of their hosts. We assessed the AMG content of viral contigs from cyanolichens by aligning predicted proteins against the KoFam HMM profiles using HMMERv3.3. Annotations were considered significant if the e-value was less than or equal to $1*10^{-5}$. The KoFam HMM profiles were downloaded (March 2019 release) at ftp://ftp.genome.jp/pub/db/kofam/archives/2019-03-20/ and selected for prokaryotic pathways [51]. Only genes involved in the category ‘Metabolism’ were included because ORFs encoding a function in DNA replication and repair are not generally considered as AMGs.

In total, we retrieved almost 250 viral contigs carrying potential AMGs from various pathways (Fig. 2). The most abundant KEGG pathways were for the metabolism of carbohydrates, nucleotides, secondary metabolites, amino acids, and terpenoids/polyketides (Fig. 2; Table S7). The abundance of KEGG pathways for secondary metabolism (e.g., biosynthesis of vancomycin, enediyne antibiotics, ansamycins, streptomycin, acarbose, and validamycin) is consistent with previous metabolomic studies of *Peltigera* that found antibiotic compounds distributed throughout the thallus [52]. The most abundant KEGG Orthology terms (KOs) for viral AMGs included KOs for methionine degradation, dTDP-L-rhamnose biosynthesis, pyrimidine deoxyribonuleotide biosynthesis, photosystem II, and fatty acid biosynthesis (Table S3). Consistent with previous studies of phages of marine cyanobacteria [11], we also identified AMGs associated with photosynthesis (e.g., *psbA* and *psbD*) on nine contigs, representing 15 ORFs and 4 ORFs, respectively. However, these viral contigs could not be assigned to hosts. Among the 50 contigs assigned to *Nostoc* as a host, 12 contigs carried at least one potential AMG (Table S2). Nine of the 27 circular viral genomes contained ORFs with potential AMGs (Table S5b).

One of the most abundant AMGs in lichen samples is for the *rfb* operon (*rfbA*, *rfbB*, *rfbC*, and *rfbD*), which is involved in the anabolism of dTDP-L-rhamnose from D-glucose-1-phosphate that is required for the biosynthesis of the O antigen of lipopolysaccharide and production of extracellular polymeric substances (EPS) in gram-negative bacteria [53]. The *rfb* operon was found on 21 viral contigs (including 1 potential prophage) in eight cyanolichen samples. Consistent with the importance of rfb for gram-negative bacteria, the majority of viral contigs with the *rfb* operon were predicted to be phages of Proteobacteria. Most contigs carried at least two of *rfb* genes: six contigs contained *rfbA* and *rfbB*; eight contigs contained *rfbA*, *rfbB* and *rfbC*; and six contained the four genes together. One contig carried only *rfbB*. The high number of contigs carrying the complete operon (identified in most cyanolichen samples) is consistent with the potential ecological importance of EPS for lichen thalli formation, water retention, and microbial communication (reviewed by [54]).

**References (not found in main text)**

21. Chagnon P, Magain N, Miadlikowska J, Lutzoni F. Species diversification and phylogenetically constrained symbiont switching generated high modularity in the lichen genus *Peltigera*. *J Ecol* 2019; **107**: 1645–1661.

22. Carbone I, White JB, Miadlikowska J, Arnold AE, Miller MA, Magain N, et al. T-BAS Version 2.1: Tree-Based Alignment Selector Toolkit for Evolutionary Placement of DNA Sequences and Viewing Alignments and Specimen Metadata on Curated and Custom Trees. *Microbiol*

*Resour Announc* 2019; **8**.

23. Carbone I, White JB, Miadlikowska J, Arnold AE, Miller MA, Kauff F, et al. T-BAS: Tree-Based Alignment Selector toolkit for phylogenetic-based placement, alignment downloads and metadata visualization: an example with the Pezizomycotina tree of life. *Bioinformatics* 2017; **33**: 1160–1168.
24. Krueger F. Trim Galore: a wrapper tool around Cutadapt and FastQC to consistently apply quality and adapter trimming to FastQ files, with some extra functionality for MspI-digested RRBS-type (Reduced Representation Bisufite-Seq) libraries. *URL http://www bioinformatics babraham ac uk/projects/trim_galore/ (Date of access: 28/04/2016)* 2012.
25. Li D, Liu C-M, Luo R, Sadakane K, Lam T-W. MEGAHIT: an ultra-fast single-node solution for large and complex metagenomics assembly via succinct de Bruijn graph. *Bioinformatics* 2015; **31**: 1674–1676.
26. Li D, Luo R, Liu C-M, Leung C-M, Ting H-F, Sadakane K, et al. MEGAHIT v1.0: A fast and scalable metagenome assembler driven by advanced methodologies and community practices. *Methods* 2016; **102**: 3–11.
27. Rikkinen J. Cyanolichens: An Evolutionary Overview. In: Rai AN, Bergman B, Rasmussen U (eds). *Cyanobacteria in Symbiosis*. 2002. Springer Netherlands, Dordrecht, pp 31–72.
28. Ren J, Ahlgren NA, Lu YY, Fuhrman JA, Sun F. VirFinder: a novel k-mer based tool for identifying viral sequences from assembled metagenomic data. *Microbiome* 2017; **5**: 69.
29. Pruitt KD, Tatusova T, Maglott DR. NCBI reference sequences (RefSeq): a curated non-redundant sequence database of genomes, transcripts and proteins. *Nucleic Acids Res* 2007; **35**: D61–5.
30. Wattam AR, Abraham D, Dalay O, Disz TL, Driscoll T, Gabbard JL, et al. PATRIC, the bacterial bioinformatics database and analysis resource. *Nucleic Acids Res* 2014; **42**: D581–91.
31. Kang DD, Li F, Kirton E, Thomas A, Egan R, An H, et al. MetaBAT 2: an adaptive binning

algorithm for robust and efficient genome reconstruction from metagenome assemblies. *PeerJ* 2019; **7**: e7359.

32. Pongor LS, Vera R, Ligeti B. Fast and sensitive alignment of microbial whole genome sequencing reads to large sequence datasets on a desktop PC: application to metagenomic datasets and pathogen identification. *PLoS One* 2014; **9**: e103441.
33. Bushnell B. BBMap: A fast, accurate, splice-aware aligner. 2014. Lawrence Berkeley National Lab. (LBNL), Berkeley, CA (United States).
34. Li H, Handsaker B, Wysoker A, Fennell T, Ruan J, Homer N, et al. The Sequence Alignment/Map format and SAMtools. *Bioinformatics* 2009; **25**: 2078–2079.
35. Hyatt D, Chen GL, LoCascio PF, Land ML, Larimer FW, Hauser LJ. Prodigal: prokaryotic gene recognition and translation initiation site identification. *BMC Bioinformatics* 2010; **11**: 119.
36. Grazziotin AL, Koonin EV, Kristensen DM. Prokaryotic Virus Orthologous Groups (pVOGs): a resource for comparative genomics and protein family annotation. *Nucleic Acids Res* 2017; **45**: D491–D498.
37. Paez-Espino D, Roux S, Chen I-MA, Palaniappan K, Ratner A, Chu K, et al. IMG/VR v.2.0: an integrated data management and analysis system for cultivated and environmental viral genomes. *Nucleic Acids Res* 2019; **47**: D678–D686.
38. Altschul SF, Gish W, Miller W, Myers EW, Lipman DJ. Basic local alignment search tool. *J Mol Biol* 1990; **215**: 403–410.
39. Gagunashvili AN, Andrésson ÓS. Distinctive characters of Nostoc genomes in cyanolichens. *BMC Genomics* 2018; **19**: 434.
40. Demovir. Github. https://github.com/feargalr/Demovir
41. Zhang F, Zhou F, Gan R, Ren C, Jia Y, Yu L, et al. PHISDetector: a tool to detect diverse in silico phage-host interaction signals for virome studies. *bioRxiv* 2020; 661074
42. Huson DH, Mitra S, Ruscheweyh H-J, Weber N, Schuster SC. Integrative analysis of

environmental sequences using MEGAN4. *Genome Res* 2011; **21**: 1552–1560.

### Supplemental Tables

**Table S1.** Mycobiont species, photobiont composition, site of collection and voucher information for 11 cyanolichen metagenomes.

| Sample ID | SRA ID | Fungal Mycobiont | Photobiont | Collection and Voucher information |
|---|---|---|---|---|
| NMS7 | SRX8034513 | *Solorina crocea* | *Coccomyxa* sp. and *Nostoc* sp. | Finland, A. Simon 79, LG, P6110 |
| JL23 | SRX8034517 | *Peltigera aphthosa* | *Coccomyxa* sp. and *Nostoc* sp. | Canada, Alberta, Miadlikowska & Lutzoni s.n., DUKE 0383963, P6003 |
| NMS8 | SRX8034514 | *Peltigera dolichorhiza* | *Nostoc* sp. | Panama, N. Magain NM23, DUKE 357954, P6117 |
| JL34 | SRX8034511 | *Peltigera evansiana* | *Nostoc* sp. | USA, Michigan, Miadlikowska & Lutzoni s.n., DUKE 0383961, P6005 |
| JL31 | SRX8034518 | *Peltigera extenuata* | *Nostoc* sp. | Canada, Alberta, Miadlikowska & Lutzoni s.n., DUKE 0383965, P6001 |
| NMS1 | SRX8034509 | *Peltigera hydrothyria* | *Nostoc* sp. | Canada, Nova Scotia, F. Anderson 16529, NSPM, P6101 |
| NMS5 | TBD | *Peltigera leucophlebia* | *Coccomyxa* sp. and *Nostoc* sp. | Iceland, N. Magain, LG, P6106 |
| JL33 | SRR11456913 | *Peltigera malacea* | *Nostoc* sp. | Canada, Alberta, Miadlikowska & Lutzoni s.n., DUKE 0383962, P6004 |
| NMS9 | SRX8034515 | *Peltigera phyllidiosa* | *Nostoc* sp. | USA, North Carolina, N. Magain, DUKE 0383964, P3103 |
| NMS3 | TBD | *Peltigera venosa* | *Coccomyxa* sp. and *Nostoc* sp. | Canada, Québec, J. Gagnon 26.15, QFA, P6103 |
| NMS4 | SRX8034512 | *Peltigera venosa* | *Coccomyxa* sp. and *Nostoc* sp. | Finland, A. Simon 80, LG, P6104 |

**Table S2**: List of AMGs and metabolic pathways identified from 12 contigs assigned to *Nostoc* as a host (out of 50 total contigs assigned to *Nostoc*).

| Sample | Contig ID | KEGG Orthology (KO) | Number of CDS | Gene | KEGG Module |
|---|---|---|---|---|---|
| JL31 | S10-k119_109292 | K00059 | 28 | *fabG* | Fatty acid biosynthesis |
| JL31 | S10-k119_1519 | K00059 | 6 | *fabG* | Fatty acid biosynthesis |
| JL31 | S1-k141_219310 | K00558 | 1 | *DNMT1* | Cysteine and methionine metabolism |
| JL23 | S11-k119_124877 | K00558 | 2 | *DNMT1* | Cysteine and methionine metabolism |
| JL34 | S13-k119_368571 | K00558 | 4 | *DNMT1* | Cysteine and methionine metabolism |
| NMS7 | S7-k141_50137 | K00558 | 9 | *DNMT1* | Cysteine and methionine metabolism |
| NMS5 | S5-k141_153852 | K02289 | 1 | *cpcF* | Photosynthesis - antenna proteins |
| NMS5 | S5-k141_196277 | K02639 | 2 | *petF* | Photosynthesis - ferredoxin |
| NMS5 | S5-k141_316970 | K02706 | 4 | *psbD* | Photosynthesis |
| JL34 | S13-k119_227010 | K03273 | 7 | *gmhB* | Lipopolysaccharide biosynthesis |
| NMS5 | S5-k141_155358 | K04091 | 5 | *ssuD* | Sulfur metabolism |
| JL23 | S11-k119_126207 | K13421 | 2 | *UMPS* | Pyrimidine metabolism |
| NMS5 | S5-k141_155358 | K15554 | 6 | *ssuC* | Sulfur metabolism |
| NMS5 | S5-k141_155358 | K15555 | 7 | *ssuB* | Sulfur metabolism |

**Table S3**. Most common KEGG KO identified in cyanolichen viral contigs.

| KEGG Orthology (KO) | Name(s) | Definition | KEGG Module | Total hits | Tri-member (% total) | Bi-member (% total) |
|---|---|---|---|---|---|---|
| K00558 | *DNMT1, dcm* | DNA (cytosine-5)-methyltransferase 1 [EC:2.1.1.37] | Methionine degradation | 47 | 17 (36%) | 30 (64%) |
| K01710 | *rfbB, rmlB, rffG* | dTDP-glucose 4,6-dehydratase [EC:4.2.1.46] | dTDP-L-rhamnose biosynthesis | 21 | 15 (71%) | 5 (24%) |
| K00973 | *rfbA, rmlA, rffH* | glucose-1-phosphate thymidylyltransferase [EC:2.7.7.24] | dTDP-L-rhamnose biosynthesis | 20 | 16 (80%) | 4 (20%) |
| K00525 | *nrdA, nrdE* | ribonucleoside-diphosphate reductase alpha chain [EC:1.17.4.1] | Pyrimidine deoxyribonuleotide biosynthesis | 18 | 12 (67%) | 6 (33%) |
| K02703 | *psbA* | photosystem II P680 reaction center D1 protein [EC:1.10.3.9] | Photosystem II | 15 | 12 (80%) | 3 (20%) |
| K00059 | *fabG, OAR1* | 3-oxoacyl-[acyl-carrier protein] reductase [EC:1.1.1.100] | Fatty acid biosynthesis | 10 | 0 (0%) | 10 (100%) |
| K00067 | *rfbD, rmlD* | dTDP-4-dehydrorhamnose reductase [EC:1.1.1.133] | dTDP-L-rhamnose biosynthesis | 10 | 8 (80%) | 2 (20%) |
| K00390 | *cysH* | phosphoadenosine phosphosulfate reductase [EC:1.8.4.8 1.8.4.10] | Assimilatory sulfate reduction | 10 | 2 (20%) | 8 (80%) |
| K01711 | *gmd, GMDS* | GDPmannose 4,6-dehydratase [EC:4.2.1.47] | Carbohydrate metabolism | 10 | 8 (80%) | 2 (20%) |
| K01790 | *rfbC, rmlC* | dTDP-4-dehydrorhamnose 3,5-epimerase [EC:5.1.3.13] | dTDP-L-rhamnose biosynthesis | 10 | 9 (90%) | 1 (10%) |
| K03465 | *thyX, thy1* | thymidylate synthase (FAD) [EC:2.1.1.148] | Pyrimidine deoxyribonuleotide biosynthesis | 10 | 3 (30%) | 7 (70%) |

The file *Ponsero_etal_SupplementalTablesS4-S8.xlsx* contains the following additional tables.

**Table S4.** List of genomes used to construct the synthetic lichen mock community.

**Table S5.** (a) Sample and functional information for 27 circular viral contigs assembled from cyanolichen metagenomes; (b) Auxiliary Metabolic Genes (AMGs) identified on circular viral genomes.

**Table S6.** Sample, contig, and taxonomic information for 1,301 cyanolichen viral contigs predicted with Marvel, Vibrant, and VirSorter.

**Table S7.** KEGG annotation for 550 AMGs (represented 904 KEGG annotations) identified on 249 cyanolichen viral contigs.

**Table S8.** Prophage sequences (n = 54) identified in *Nostoc* genomes with VirSorter.

### Supplementary Figures and Legends

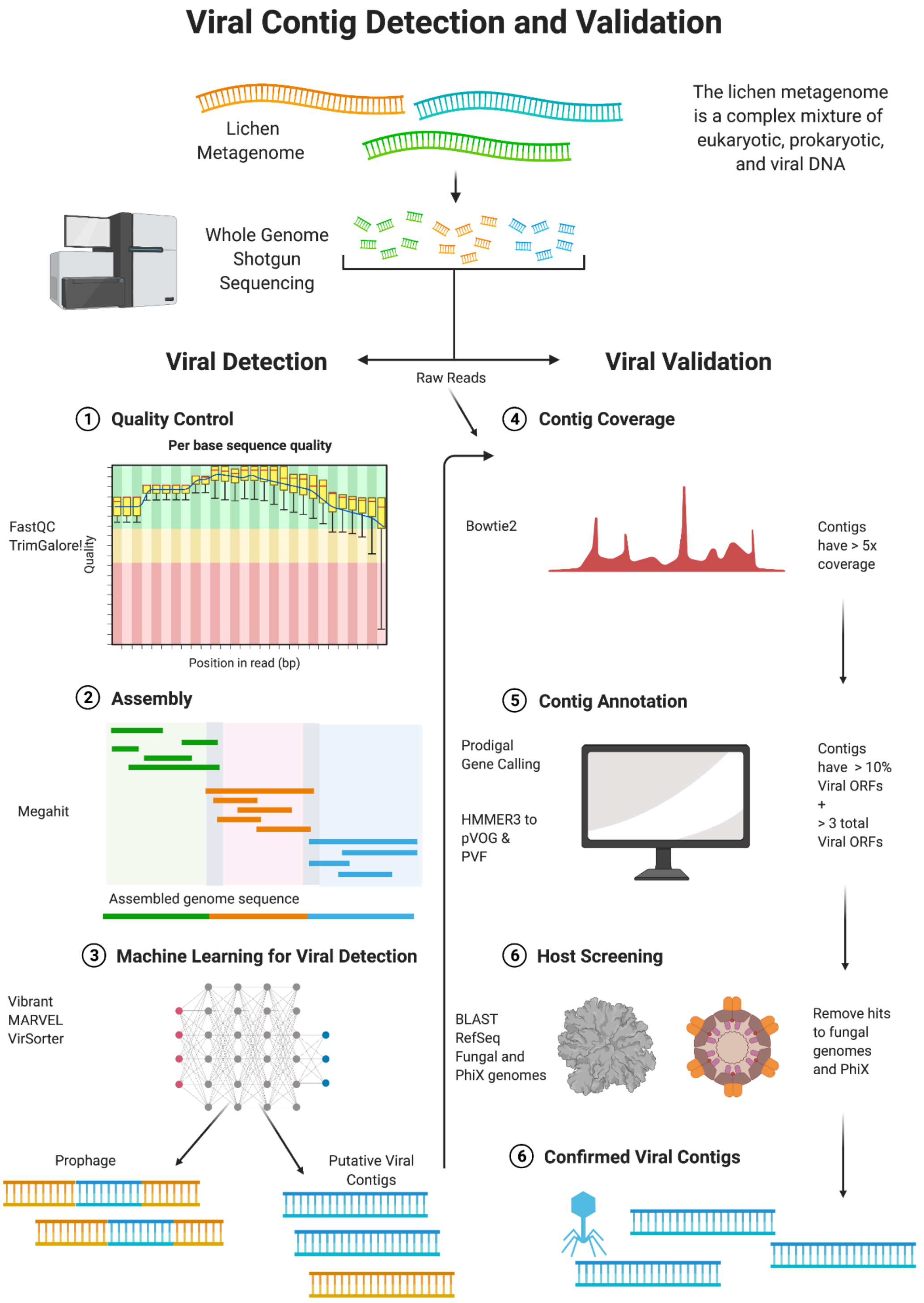


**Figure S1:** Analytical workflow for the retrieval of viral sequences from cyanolichen metagenomes.

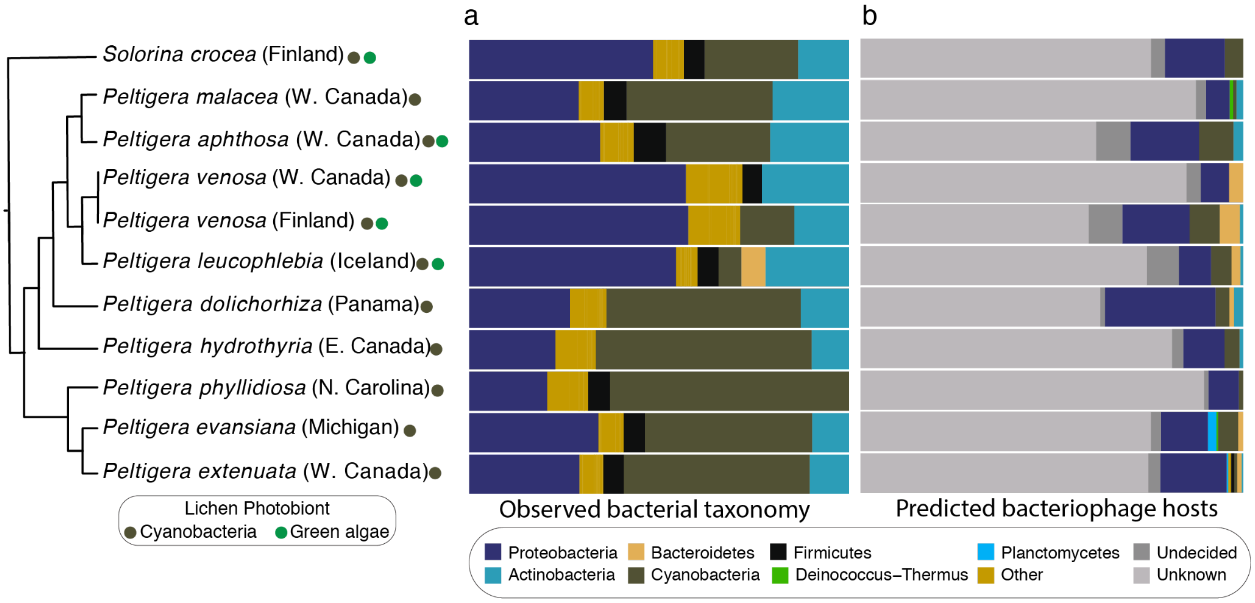


**Figure S2. Congruence between bacterial taxonomy and predicted cyanolichen viral host range. (a)** Phylum level taxonomy of bacterial sequences in cyanolichen metagenomes; and **(b)** viral host taxonomy as predicted with Demovir [40]. Phylogenetic tree follows Fig. 1.

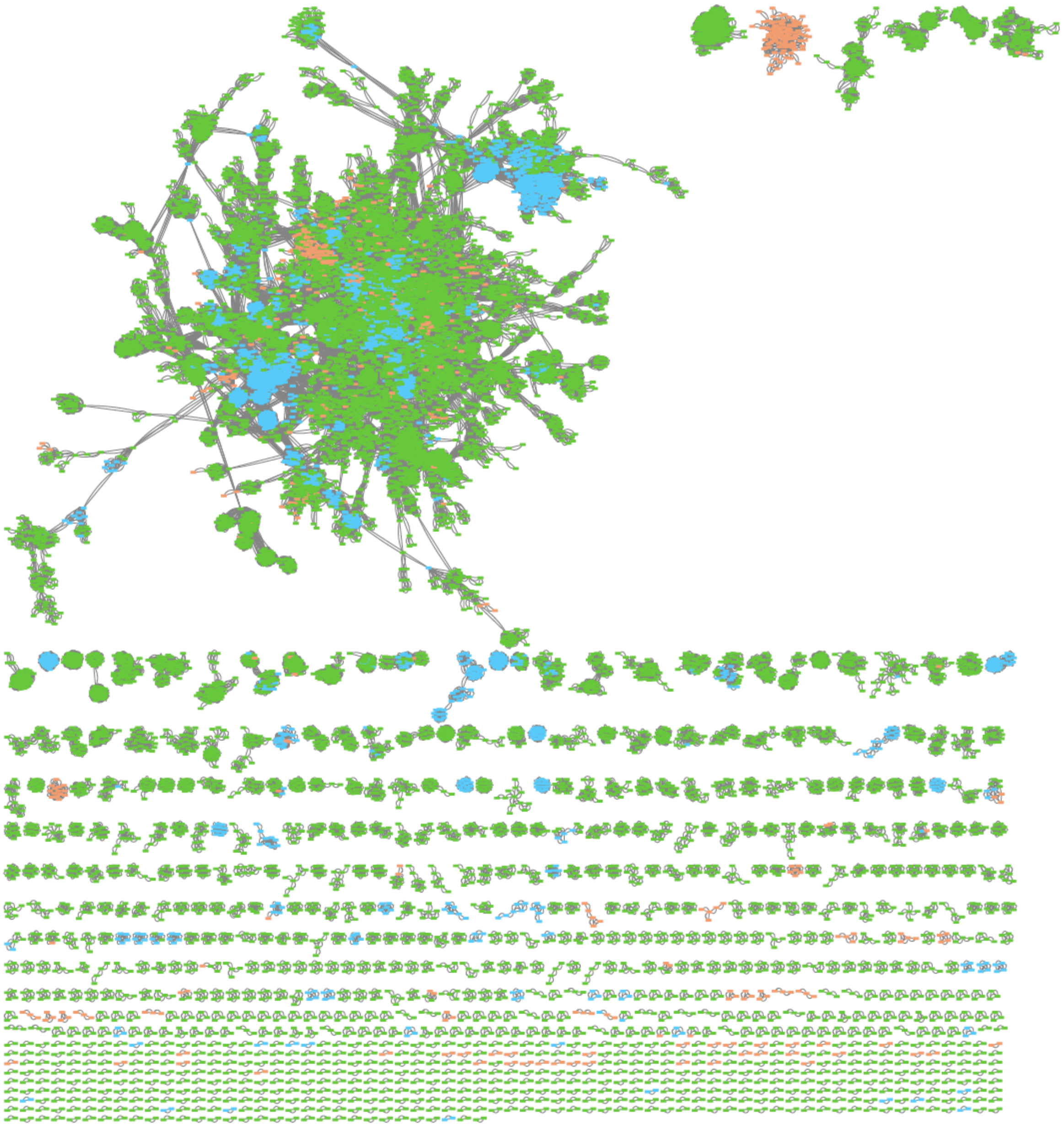

**Figure S3: Network of viral contigs (VCs) illustrates uniqueness of cyanlichen VCs.** Colors indicate data source (orange: viral contigs from cyanolichen metagenomes; blue: RefSeq viral genomes; and green: viral contigs from IMG/VR representing other terrestrial habitats, such as "Soil", "Rhizoplane", "Rhizosphere", "Phyllosphere" and "Plant litter").

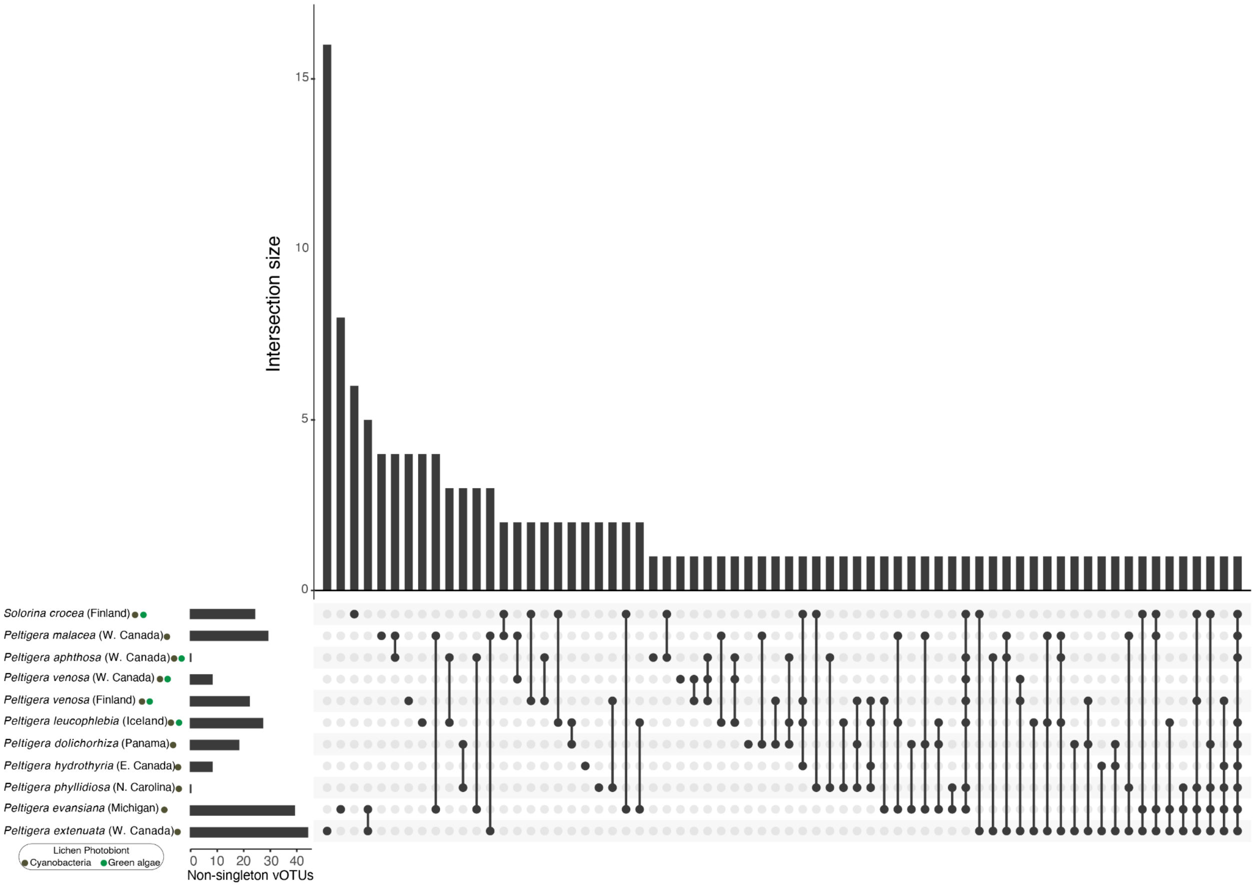


**Figure S4**: **Cyanolichen metagenomes contain largely distinct viral clusters (VCs)**. UpSet Plot illustrates little overlap in VCs among 11 metagenome samples. Bi- and tri-membered lichen thalli shared 47 VCs (57 and 29 were unique to bi- or tri-membered thalli, respectively). No VCs were shared among all cyanolichen samples.

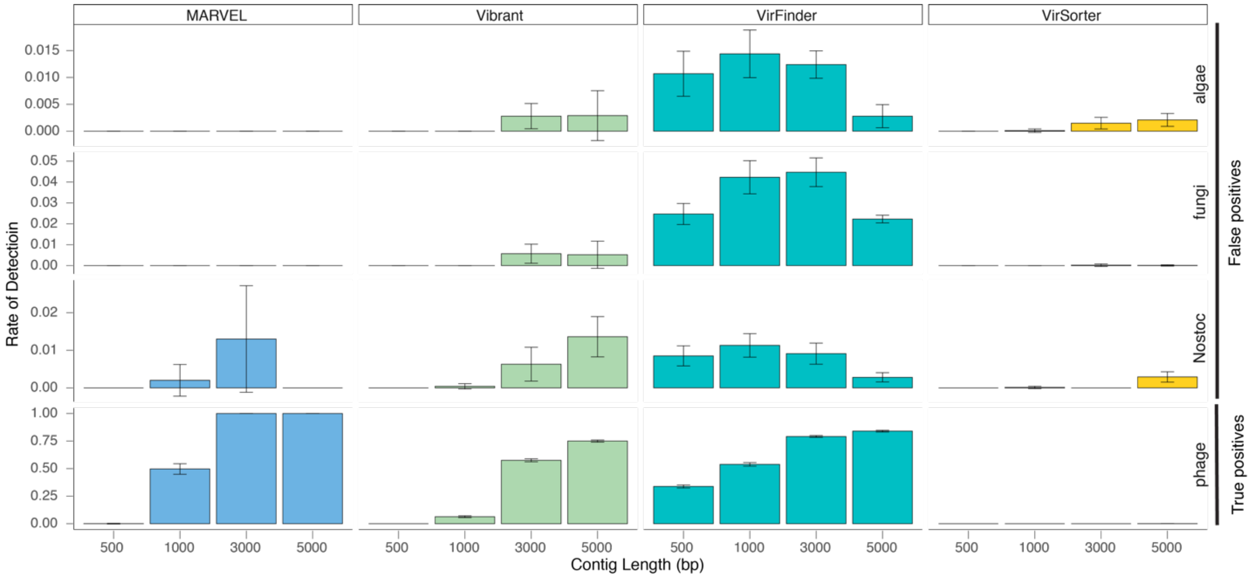


**Figure S5: Performance comparison of MARVEL, Vibrant, VirSorter, and VirFinder on synthetic lichen mock communities.** Error bars show the standard deviation of 10 independent replicates of the synthetic mock community. The top three rows represent the rate of false positives (i.e., algal, fungal, or *Nostoc* sequences that were falsely identified as being viral), whereas the bottom row demonstrates the rate of detection for true positives (i.e., phage sequences).

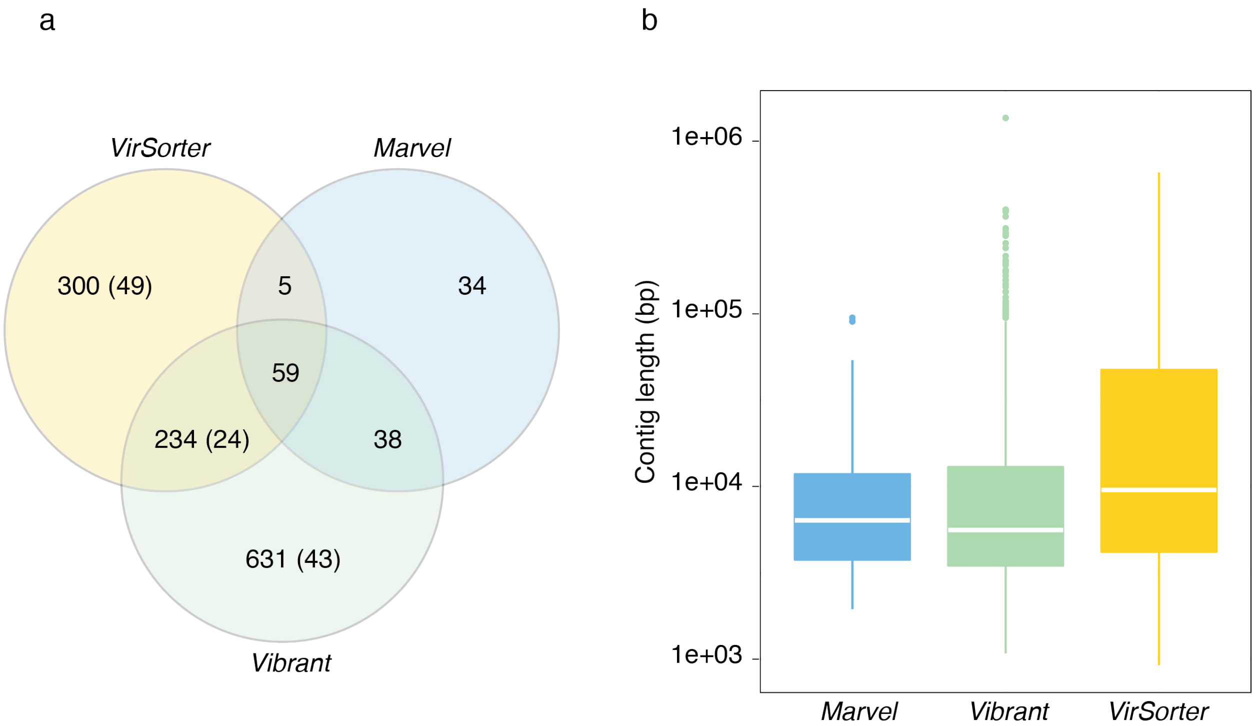


**Figure S6. Overlap and length distribution of high-quality viral contigs identified with different viral detection methods.** (**a**) Venn diagram predicted viral contigs for three different machine-learning viral detection tools. Number of predicted prophages for VirSorter and Vibrant are shown in parentheses. (**b**) Box plot with median length and interquartile distribution of viral contigs retrieved by each method.
